# Heterogeneity of ictal firing during generalized seizures in the awake cortex

**DOI:** 10.1101/2025.10.14.682021

**Authors:** Péter Sere, Nikolett Zsigri, Vincenzo Crunelli, Magor L. Lőrincz

## Abstract

Cortico-thalamo-cortical oscillations are central to both normal and pathological brain activities and emerge from complex cortical and thalamic interactions. However, the specific activity of identified cortical neurons during the paroxysmal oscillations associated with absence seizures (ASs) in awake animals remains underexplored. The dominant narrative suggests that seizures indiscriminately disrupt cortical activity through generalized hyperexcitability, but direct evidence supporting this view is lacking. Here, we recorded single units from pyramidal neurons and different interneuron subtypes in the neocortex of two validated rodent models of absence epilepsy under awake, behaving conditions. We find that neurons maintain their firing rank order across interictal and ictal states, regardless of whether their ictal firing rate increases, decreases, or remains stable compared to the interictal phase. Rather than a random cortical takeover, ictal activity represents a scalable modulation of pre-existing network states. These results challenge the generalized hyperexcitability model and highlight the structured, heterogeneous nature of cortical activity during ASs, with implications for mechanistic understanding and targeted therapies.

## INTRODUCTION

Epilepsy is a debilitating neurological disorder affecting approximately 1% of the global population, leading to diminished quality of life, comorbidities, and mortality (Fisher et al., 2014). Despite extensive research using diverse *in vivo* and *in vitro* models, the cellular mechanisms underlying seizure generation, propagation and termination remain incompletely understood. The prevailing view portrays cortical activity during seizures as homogeneous and hypersynchronous, driven by disrupted excitation–inhibition balance (Engel, 1996). However, mounting evidence indicates that neuronal firing is far more heterogeneous in both experimental models (Bower and Buckmaster, 2008; Matsumoto and Marsan, 1964; McCafferty et al., 2023; Meyer et al., 2018; Sawa et al., 1968) and human patients (Truccolo et al., 2011). Indeed, human single-unit studies during both interictal events and seizures reveal marked heterogeneity: the firing rate of only a subset of neurons is modulated during interictal discharges with diverse patterns—including pre-event increases and decreases near the focus—and seizure initiation/spread is characterized by mixed rate changes rather than monolithic recruitment (Keller et al., 2010; Truccolo et al., 2011).

The cerebral cortex hosts a diverse population of neurons that differ in morphology (Szentágothai, 1978), electrophysiology (McCormick et al., 1985) and neurochemistry (Lee et al., 2010). Recent transcriptomic studies have revealed even greater diversity (Gouwens et al., 2020), with neuronal identity linked more closely to brain state than to morphology or intrinsic physiology (Bugeon et al., 2022). These findings suggest that structured neuronal heterogeneity might be a defining feature of cortical organization. Indeed, during physiological oscillations such as sleep slow waves (Steriade et al., 1993), gamma (Varga et al., 2014), alpha (Lorincz et al., 2009), or sleep spindle (Gardner et al., 2013) oscillations neurons exhibit variable firing across cycles. Thus, synchronized network rhythms can coexist with pronounced single-cell variability—implying that heterogeneity is not noise, but an organized property of cortical dynamics.

Paroxysmal activity, characterized by excessive synchrony and large-amplitude EEG deflections, is generally attributed to altered excitation/inhibition balance or hyperexcitability (Lőrincz et al., 2024). Work *in vitro* and anesthetized preparations suggest that cortical neurons discharge synchronously during such events (Ziburkus et al., 2006). Yet, studies in awake behaving animals have revealed a different picture: cortical (McCafferty et al., 2023; Meyer et al., 2018), thalamic (McCafferty et al., 2023, 2018), and striatal (Miyamoto et al., 2019) neurons show variable and less entrained activity. This discrepancy suggests that anesthesia may artificially enhance synchrony, masking the intrinsic heterogeneity present during seizures under natural conditions.

Thus, the neuronal dynamics underlying seizure generation, propagation, and termination in the awake cortex remain poorly understood. The dominant narrative suggests that seizures overtake cortical networks in a chaotic and indiscriminate manner, but the supportive evidence is lacking. If physiological brain rhythms already express structured heterogeneity, could pathological oscillations such as spike-and-wave discharges (SWDs) of absence seizures instead reflect a scalable modulation of pre-existing network states rather than a random cortical takeover? Building on ensemble studies that documented mixed ictal rate changes in cortex and thalamus in awake rodents (McCafferty et al., 2023, 2018; Meyer et al., 2018; Miyamoto et al., 2019), we provide cell-type–resolved, single-neuron recordings from identified pyramidal, regular-spiking, and fast-spiking neurons in awake cortex across two validated models of absence epilepsy. We find that neurons maintain their firing rank order across interictal and ictal states, regardless of whether their ictal firing rate increases, decreases, or remains stable compared to the interictal state. Moreover, the magnitude of ictal modulation inversely scales with baseline rate and covaries with spike-and-wave entrainment, revealing a principled structure to this heterogeneity. Thus, SWDs emerge as a scalable reweighting of cortical dynamics—extending prior work with cell-type and cross-species resolution—and challenge the hyperexcitability trope, indicating a structured, hierarchical modulation of cortical networks.

## Materials and Methods

All procedures complied with the European Communities Council Directives of 1986 (86/609/EEC) and 2003 (2003/65/CE) for animal research and were approved by the Ethics Committee of the University of Szeged. We used Stargazer mice (n = 5, 2 females; 3–7 months of age), a well-established monogenic model of absence epilepsy (Noebels et al., 1990), and Genetic Absence Epilepsy Rats from Strasbourg (GAERS) (n = 7, 4 females; 3–7 months), a validated polygenic model (Depaulis et al., 2016). Animals were maintained on a 12:12 h light:dark cycle with ad libitum access to food and water. They were group-housed prior to surgery and singly housed thereafter to protect head implants.

Habituation and recording procedures followed our previous work (McCafferty et al., 2023; Molnár et al., 2021). Briefly, anesthesia was induced with 4% isoflurane in oxygen (2 L/min) in an induction chamber and maintained via a facemask with active scavenging. During surgery, isoflurane was gradually reduced to 1.2–1.8% in oxygen (1 L/min), with depth monitored by respiratory rate and pedal reflex. Animals were mounted in a stereotaxic frame (Model 902, David Kopf Instruments, USA), the skull was exposed, and a stainless-steel head post was cemented over the frontal suture using dental acrylic (Pattern Resin LS, GC America, USA). Craniotomy sites were marked at the following coordinates: rat S1 (primary somatosensory cortex): 0.0 mm anteroposterior, 2.5 mm lateral to bregma; mouse M1 (primary motor cortex): −1.8 mm anteroposterior, 2.0 mm lateral to bregma.

For postoperative care, animals received carprofen (Rimadyl; 5 mg/kg, i.p.; Pfizer, USA) and gentamicin (0.1 mg/kg, i.m.). After at least 5 days of recovery, animals were handled daily for 7–21 days to reduce stress during head fixation, with fixation duration increased gradually.

On recording days, small craniotomies (0.8–1 mm) were made under isoflurane (Forane, AbbVie, USA; 1.0–1.5% in OC at 1 L/min) at the marked positions, leaving the dura intact. Mineral oil was applied to prevent dural dehydration. Animals were then transferred to the in vivo electrophysiology setup; head posts were secured in a custom holder, and recording began ≥40 min after full recovery from anesthesia.

Electrophysiology. Single-unit extracellular recordings were obtained in rat S1 and mouse M1 using either glass micropipettes filled with 0.5 M NaCl (impedance 3–8 MΩ) or single-shank, 32-channel silicon probes (NeuroNexus, USA). A nearby (≈0.3 mm) insulated wire (Wf wire; [specify material/manufacturer]) positioned in infragranular layers was used to record the local field potential (LFP). Glass-electrode signals were preamplified with Axon HS-9A headstages (Molecular Devices), amplified with an Axoclamp 900A (Molecular Devices, USA), and filtered at 0.3–6 kHz for single units. LFPs (0.1–200 Hz) were recorded with an AC amplifier (Supertech, Hungary; gain 1000×). Signals were digitized at 30 kHz using a CED Power1401-3 A/D converter (CED, UK) and Spike2 software. Silicon-probe data were amplified, filtered, and digitized with an Intan RHD2000 board (Intan Technologies) at 30 kHz. Offline analysis was performed in Spike2 (CED, UK) and OriginPro 8.5 (Microcal, USA); figures were prepared in OriginPro 8.5. Data are reported as mean ± SD unless noted.

Seizure detection. Seizure times were first identified by amplitude threshold crossings in the LFP and then confirmed by the presence of spike-and-wave discharges (SWDs) at 6–8 Hz with a minimum duration of ≥3 s.

The modulation index (MI) was computed as

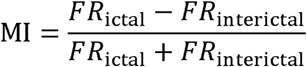

such that MI = +1 denotes maximal enhancement and MI = −1 denotes maximal suppression of firing during the interictal→ictal transition. For each neuron and state (interictal, ictal), spiking regularity was quantified as the coefficient of variation of interspike intervals: CV = σISI/μISI. ISIs were computed from all spikes within a state; neurons contributing fewer than 20 spikes in a state were excluded from CV analyses. The entrainment index (EI) was defined as the fraction of spikes occurring while the SWD “spike” component of each cycle exceeded 50% of its peak amplitude. The rhythmicity index (RI) was defined as the amplitude ratio of the first side-peak to the main peak in the neuron’s ictal spike-train autocorrelogram.

Unless otherwise stated, the experimental unit was the animal. When multiple neurons were recorded from the same animal, single-cell measures (e.g., firing rate, MI, EI, RI, CV) were first aggregated per animal to avoid pseudo-replication; group comparisons were then performed on these per-animal summaries. For each outcome, we inspected histograms and Q–Q plots of per-animal summaries and tested normality with Shapiro–Wilk. When normality and homoscedasticity were satisfied (Levene’s test), we used parametric tests; otherwise we used non-parametric alternatives. When comparing two groups we used unpaired t-test or Mann–Whitney U (Wilcoxon rank-sum). When comparing three groups (e.g., PYR vs RSI vs FSI) we used one-way ANOVA. When comparing paired states (interictal vs ictal within animals) the Wilcoxon signed-rank test was used. All tests were two-tailed with α = 0.05. Exact P values are reported. We avoided t-tests when >2 conditions were compared. Animals were assigned to surgical preparation and recording order without formal randomization; no group allocation was involved beyond species/strain. Seizure detection used predefined thresholds and frequency/duration criteria and was run blind to neuron identity. Spike sorting and unit inclusion were performed with automated criteria followed by an investigator blind to cell-type labels. Given the exploratory nature and technical constraints of awake single-unit recordings, a priori power calculations were not performed; instead we report effect sizes and confidence intervals for all key comparisons.

## RESULTS

Because ictal activity is characterized by increased neuronal synchrony that can obscure the distinction of individual units (Huguenard, 2019), we performed single-unit juxtacellular recordings in the infragranular layers of the primary somatosensory (S1) barrel cortex of awake, head-restrained GAERS rats and in the primary motor cortex (M1) of Stargazer mice. From these experiments, we obtained 52 well-isolated neurons (24 from rats, 28 from mice) that exhibited sufficient interictal and ictal epochs for analysis. The use of high signal-to-noise, single-cell recordings ensured that even interictally recorded spikes could be unambiguously attributed to individual neurons (Fig. 1A–B).

**Fig. 1.**
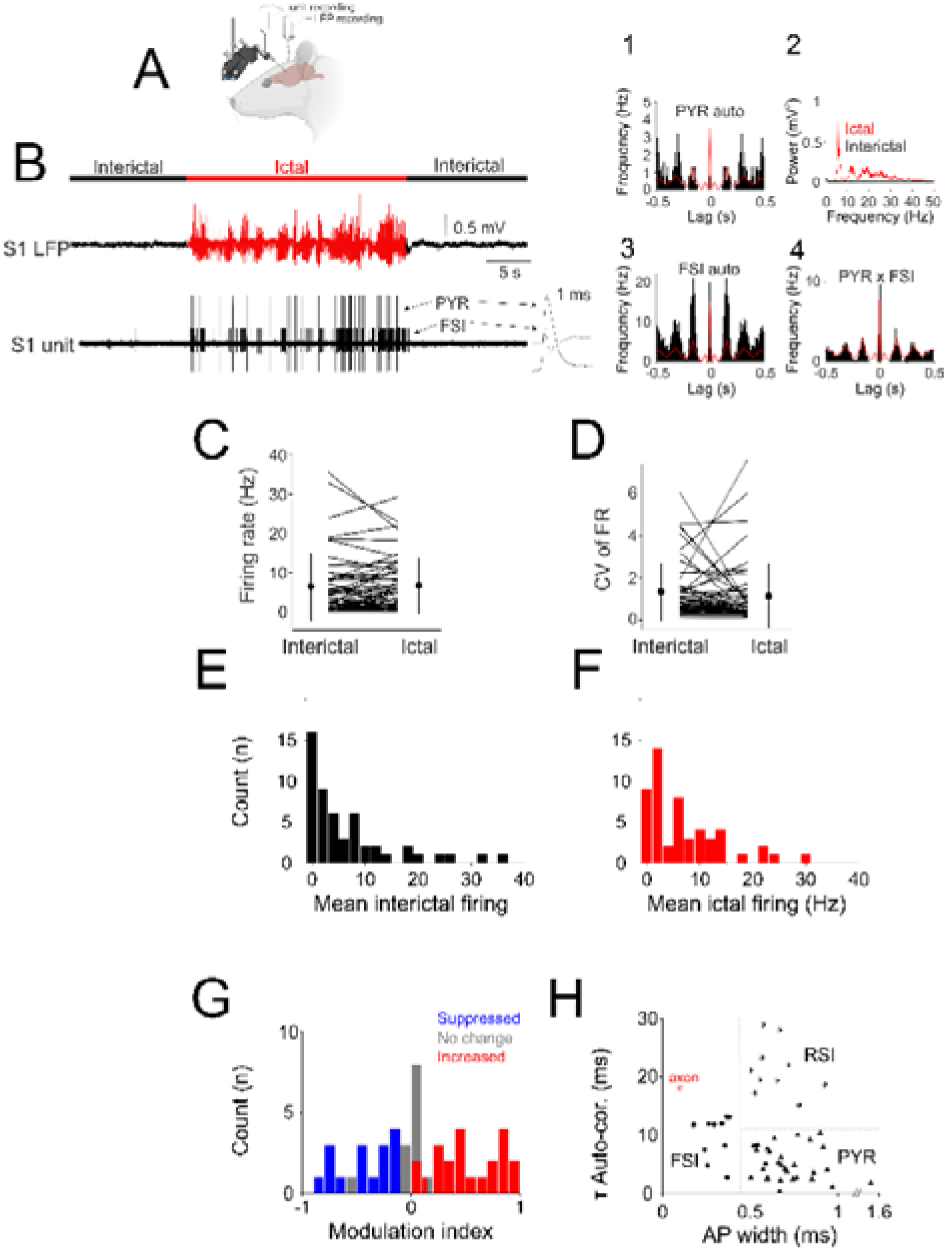
Ictal and interictal firing properties of cortical neurons. (A) Schematics of the experimental design. (B) Example simultaneous recording of two cortical neurons (PYR: pyramidal neuron; FSI: fast-spiking interneuron) characterized by ictal entrainment in the S1BF of a GAERS rat. The interictal and ictal periods are indicated with black and red traces, respectively. Averaged action potentials (mean±s.e.m) are shown on the right on a faster timebase. Ictal autocorrelations of the pyramidal neuron (B1) and fast-spiking interneuron (B2) and their cross-correlation (B3) are shown with the LFP autocorrelation overdrawn in red. Power spectral densities of the LFP interictal (black) and ictal (red) periods are shown in (B4). (C) Mean interictal and ictal firing rates of all recorded neurons. Lines indicate individual neurons; circles indicate mean and standard deviation. (D) Coefficients of variation (CVV) of interictal and ictal mean firing rates for all recorded neurons. Lines indicate individual neurons; circles indicate mean and standard deviation. (E) Distribution of mean ictal firing rates. (F) Distribution of mean interictal firing rates. (G) Distribution of modulation indices (MI). Blue bars indicate significant ictal suppression, gray bars indicate no significant change, and red bars indicate significant ictal firing rate increase, respectively. (H) Identification of the recorded neurons. Scatter plot of interictal autocorrelation time constants (τ Auto-cor.) vs. action potential width (AP width) clearly separates fast-spiking interneurons (FSI), regular-spiking interneurons (RSI), and pyramidal neurons (PYR).

Across all recorded neurons, mean interictal (6.61 ± 8.43 Hz) and ictal (6.80 ± 6.95 Hz) firing rates did not differ significantly (p = 0.394, Wilcoxon rank-sum test, n = 52, Fig. 1C). Similarly, the coefficients of variation (CVs) of interictal and ictal firing were comparable (1.32 ± 1.34 vs. 1.12 ± 1.47, p = 0.099; Fig. 1D). Nevertheless, interictal and ictal firing rates were strongly correlated (ρ = 0.780, p = 9.6 × 10⁻¹²), indicating that neurons retained their relative firing hierarchy across brain states. To quantify cell-specific changes, we calculated a modulation index (MI; –1 = maximal suppression, +1 = maximal increase). The MI distribution revealed pronounced heterogeneity, with approximately three-quarters of neurons (39/52, 75%) showing significant modulation between interictal and ictal periods (p < 0.05, Wilcoxon signed-rank test for individual neurons). Among these, 22 neurons (42%) increased their firing, 17 (33%) decreased it, and 13 (25%) showed no significant rate change (Fig. 1E–G). The absolute MI was negatively correlated with baseline interictal firing (r = –0.459, p = 6.3 × 10⁻⁴), suggesting that neurons with higher baseline activity were less likely to undergo strong ictal modulation.

Neurons were classified as pyramidal (PYR), regular-spiking interneurons (RSI), or fast-spiking interneurons (FSI) based on spike waveform width and autocorrelation time constant (τ) following established cteria (Petersen et al., 2021). In GAERS rats, we identified 13 PYR, 5 RSI, and 6 FSI neurons, while in Stargazer mice the distribution was similar, with 19 PYR, 5 RSI, and 4 FSI neurons. The electrophysiological properties of corresponding cell types were comparable across species (p > 0.19 for PYR spike width and τ; p = 0.26 for FSI τ; p = 1.0 for RSI width), supporting pooled analyses (Fig. 1H). Thus, the two datasets were sufficiently similar to examine ictal modulation at both population and cell-type levels.

Across all neurons, FSIs fired faster than PYR and RSI neurons during interictal periods (FSI: 14.82 ± 12.49 Hz; PYR: 5.88 ± 9.86 Hz; RSI: 5.76 ± 6.94 Hz; Kruskal–Wallis p =

0.0097), but during seizures mean firing rates converged across types (p = 0.70). Firing was temporally more regular in FSIs (ictal CV = 0.54 ± 0.77) than in PYR (1.09 ± 1.25) or RSI (1.74 ± 2.34; p = 0.046). Neither the MI nor the entrainment index (EI) differed significantly across cell types (p ≥ 0.70). Within each group, ictal rate modulation was heterogeneous. Among pyramidal neurons, 15 (48%) increased, 8 (26%) decreased, and 8 (26%) showed no significant change in firing rate (interictal 4.42 ± 5.55 Hz; ictal 4.71 ± 4.36 Hz; p = 0.20; Fig. 2). Among regular-spiking interneurons, 3 (30%) increased, 5 (50%) decreased, and 2 (20%) remained unchanged (interictal 5.76 ± 6.94 Hz; ictal 6.24 ± 8.82 Hz; p = 0.85; Fig. 3). Fast-spiking interneurons showed a similar pattern, with 3 (30%) increasing, 4 (40%) decreasing, and 3 (30%) showing no significant change (interictal 14.82 ± 12.49 Hz; ictal 14.31 ± 7.19 Hz; p = 1.0; Fig. 4). These proportions were comparable in GAERS and Stargazer datasets, indicating that heterogeneous modulation characterized all major cortical cell types independent of species or cortical area.

**Fig. 2.**
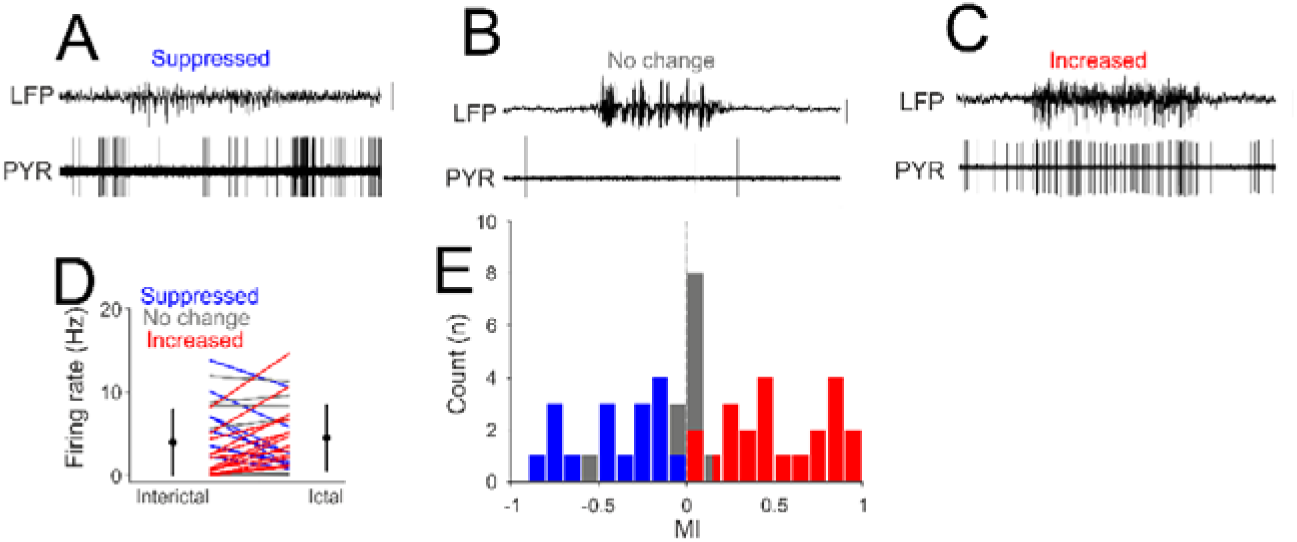
Ictal and interictal firing properties of neocortical pyramidal neurons. (A) Example simultaneous recording of M1 LFP and single unit recording of a pyramidal neuron of a Stargazer mouse characterized by suppressed ictal firing activity. (B) Example simultaneous recording of S1BF LFP and single unit recording of a pyramidal neuron characterized by unaltered ictal firing activity. (C) Example simultaneous recording of S1BF LFP and single unit recording of a pyramidal neuron characterized by enhanced ictal firing activity. (D) Mean interictal and ictal firing rates of all pyramidal neurons recorded. Lines indicate individual neurons; circles indicate mean and standard deviation. Blue lines and bars indicate significant ictal suppression, gray lines and bars indicate no significant change, and red lines and bars indicate significant ictal increase in firing rates, respectively. (E) Distribution of modulation indexes (MI) in all recorded pyramidal neurons. Calibrations: 0.5 mV, 5 seconds.

**Fig. 3.**
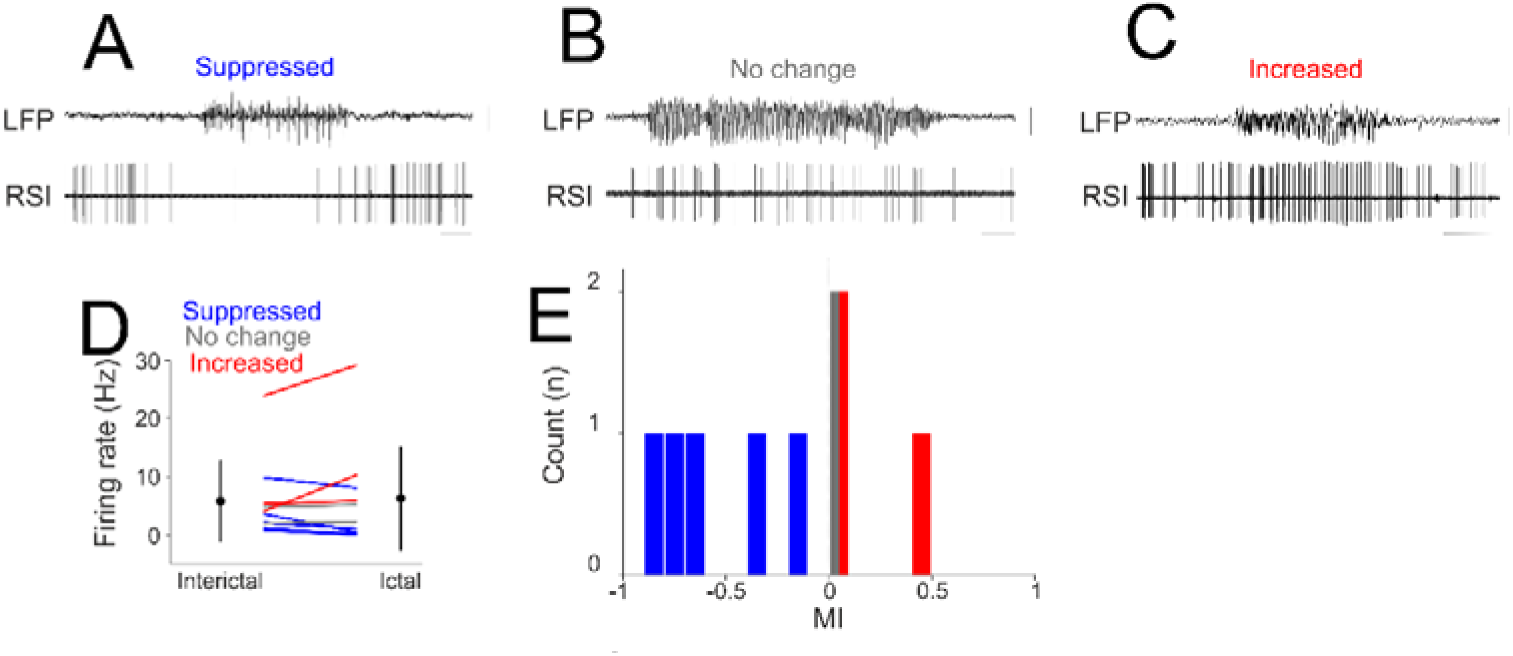
Ictal and interictal firing properties of neocortical regular spiking interneurons. Example simultaneous recording of S1BF LFP and single unit recording of a regular spiking interneuron characterized by suppressed ictal firing activity. (B) Example simultaneous recording of S1BF LFP and single unit recording of a regular spiking interneuron characterized by unaltered ictal firing activity. (C) Example simultaneous recording of S1BF LFP and single unit recording of a regular spiking interneuron characterized by enhanced ictal firing activity. (D) Mean interictal and ictal firing rates of all pyramidal neurons recorded. Lines indicate individual neurons; circles indicate mean and standard deviation. Blue lines and bars indicate significant ictal suppression, gray lines and bars indicate no significant change and red lines and bars indicate significant ictal increase in firing rates, respectively. (E) Distribution of modulation indexes (MI) in all recorded FSIs. Calibrations: 0.5 mV, 5 seconds.

**Fig. 4.**
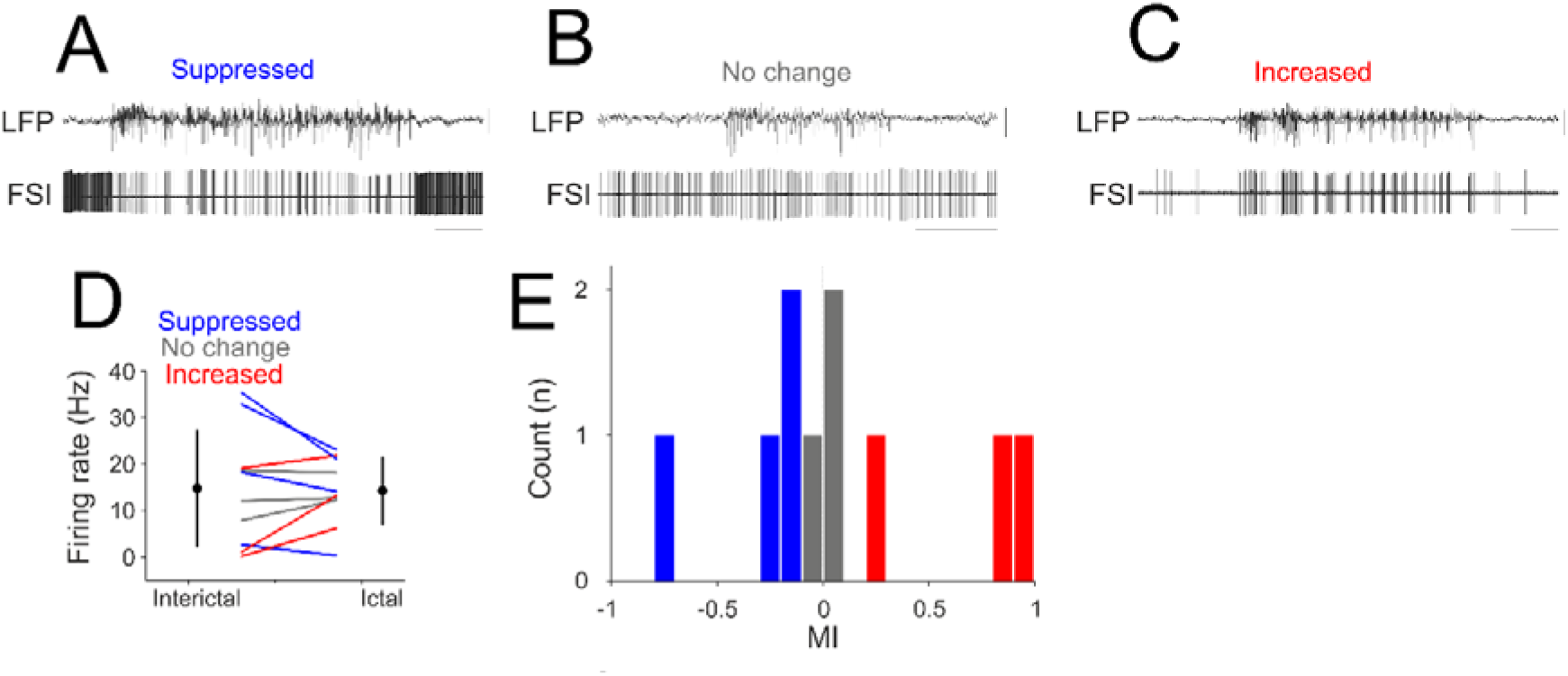
Ictal and interictal firing properties of neocortical fast-spiking interneurons. Example simultaneous recording of S1BF LFP and single unit recording of a fast-spiking interneuron characterized by suppressed ictal firing activity. (B) Example simultaneous recording of S1BF LFP and single unit recording of a fast-spiking interneuron characterized by unaltered ictal firing activity. (C) Example simultaneous recording of S1BF LFP and single unit recording of a fast-spiking interneuron characterized by enhanced ictal firing activity. (D) Mean interictal and ictal firing rates of FSIs recorded. Lines indicate individual neurons; circles indicate mean and standard deviation. Blue lines and bars indicate significant ictal suppression, gray lines and bars indicate no significant change and red lines and bars indicate significant ictal increase in firing rates, respectively. (E) Distribution of modulation indexes (MI) in all recorded FSIs. Calibrations: 0.5 mV, 5 seconds.

Across the entire population, interictal and ictal firing rates remained strongly correlated (r = 0.79, p = 1.6 × 10⁻¹²; Fig. 5A), and firing variability (CV) was likewise preserved across states (r = 0.67, p = 4.6 × 10⁻⁸; data not illustrated). The modulation index (MI) was inversely related to baseline rate (r = –0.44, p = 0.0011; Fig. 5B), confirming that low-firing neurons tended to show stronger ictal changes. Thus, neuronal heterogeneity was not random but scaled with intrinsic excitability—high-rate neurons remained stable, whereas low-rate neurons exhibited the largest ictal modulation.

**Fig. 5.**
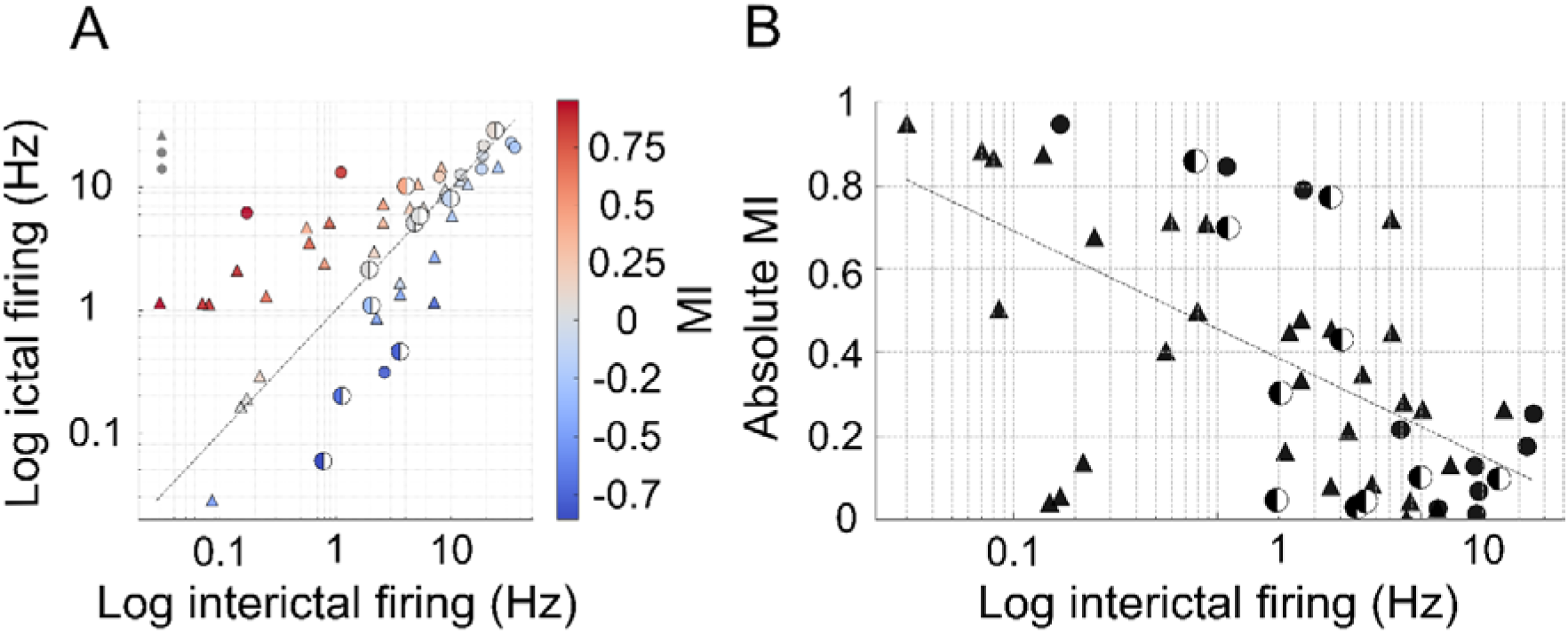
Firing rate modulation of neocortical neurons during absence seizures. (A) Scatter plot of interictal vs. ictal firing rates (log–log scale). Each point represents a single neuron, with symbol shape indicating cell type (▴ pyramidal neurons, ● fast-spiking interneurons, Ill regular-spiking interneurons). Colors denote modulation index (MI), with warmer hues indicating ictal increases and cooler hues indicating decreases. The dashed unity line indicates equal ictal and interictal firing. (B) Relationship between baseline interictal firing rate and the absolute modulation index (MI). Symbols and colors follow the same convention as in (A). The dashed line shows a log-linear regression fit, illustrating the negative correlation between baseline rate and modulation strength.

We next examined the temporal structure of firing using two complementary metrics. The rhythmicity index (RI), defined as the ratio of the first side-peak to the mean peak in autocorrelations, captured cycle-to-cycle regularity, whereas the entrainment index (EI) quantified the fraction of spikes occurring during the LFP “spike” phase of spike-and-wave discharges (SWDs). RI values averaged 0.88 ± 0.58 for PYR, 1.25 ± 0.60 for RSI, and 0.77 ±0.28 for FSI, with no significant group differences (p > 0.05, one-way ANOVA). RI did not correlate with either MI (r = 0.11, p = 0.61) or ictal firing rate. In contrast, EI was positively correlated with MI (r = 0.39, p = 0.0043; data not illustrated), indicating that neurons more strongly phase-locked to SWD cycles also exhibited greater firing-rate modulation. These analyses reveal that ictal modulation comprises two distinct but related dimensions: overall rate scaling (MI) and temporal alignment (EI). The coexistence of stable firing rank order with graded entrainment demonstrates that seizures reorganize, rather than randomize, cortical activity.

Across cell types, cortical neurons in awake models of absence epilepsy exhibited markedly heterogeneous ictal responses. Despite wide variation in the direction and magnitude of firing changes, neurons preserved their relative firing hierarchy and firing variability across brain states. The strength of ictal modulation scaled inversely with baseline activity and correlated with spike-and-wave entrainment, indicating structured, cell-specific modulation rather than generalized hyperexcitability. These findings suggest that absence seizures represent a scalable reconfiguration of pre-existing cortical network dynamics rather than a random or uniform cortical takeover.

## DISCUSSION

By recording single cortical neurons in awake models of absence epilepsy, we demonstrate that neuronal activity during SWDs is highly heterogeneous yet structured. Despite substantial variability in individual firing rates, neurons preserved their relative firing rank and variability across interictal and ictal states. These results indicate that seizures do not impose an uniform excitatory drive on the cortex but instead represent a scalable modulation of pre-existing network organization. This finding challenges the classical view that seizures arise from generalized cortical hypersynchrony and supports an alternative framework in which pathological oscillations reflect reorganized but intrinsically structured network activity.

Previous studies performed under anesthesia have reported highly stereotyped rhythmic bursting of cortical, thalamic, and striatal neurons during absence seizures (Crunelli et al., 2020). In contrast, recent work in awake behaving animals has shown greater variability and weaker entrainment of neuronal firing (McCafferty et al., 2018, 2023; Meyer et al., 2018; Miyamoto et al., 2019). Our results extend these observations by demonstrating that ictal heterogeneity is not limited to population averages but is a fundamental property of individual cortical neurons, encompassing both excitatory and inhibitory cell types in the cortical initiation zone of SWDs. This suggests that the homogeneity observed in anesthetized preparations likely reflects the artificial synchrony induced by reduced brain-state dynamics rather than a defining feature of absence seizures themselves.

The preservation of firing rank order together with graded entrainment mirrors human findings where conditional-inference and maximum-entropy models attribute most fine synchrony to nonstationary rate modulation and local network activation, supporting a scaled, structured modulation rather than uniform hypersynchrony (Keller et al., 2010; Truccolo et al., 2011). Diversity at the single-unit level is present during both interictal discharges and ictal epochs in humans, with pre-event modulations localized near the focus and heterogeneous fast-component responses, a finding that parallels our awake cortical results (Keller et al., 2010; Truccolo et al., 2011).

The coexistence of heterogeneous cellular firing with stereotypical EEG SWDs implies that cortical networks employ normalization mechanisms to stabilize overall output despite diverse single-cell responses. In this framework, seizures maintain consistent macroscopic patterns by scaling the activity of individual neurons relative to the collective network drive. Similar divisive normalization processes have been described in sensory and motor cortices, where neuronal responses are proportional to the pooled activity of neighboring neurons (Carandini and Heeger, 2011). The preserved firing rank order observed here is consistent with such a normalization principle, suggesting that absence seizures modulate the amplitude, but not the structure, of cortical activity.

Several cellular and synaptic mechanisms could underlie this normalization. Conductance increases—particularly those mediated by GABAA receptor–driven shunting inhibition—can scale membrane potential responses without altering net synaptic current (Borg-Graham et al., 1998). Balanced elevations of excitatory and inhibitory conductances can therefore adjust neuronal gain divisively (Silver, 2010), enabling neurons to maintain proportional firing across brain states. Stochastic fluctuations in synaptic input from connected neurons may further induce dynamic conductance changes that reinforce this divisive scaling (Chance et al., 2002). Together, these mechanisms can yield heterogeneous single-cell firing while maintaining stable network-level activity, reconciling cellular variability with the stereotyped EEG manifestations of SWDs.

Beyond local circuit mechanisms, broader network interactions likely contribute to the observed dynamics. Infragranular layers of the barrel cortex receive substantial non-sensory input from motor regions, both directly and via the posterior-medial thalamic nucleus (Deschênes et al., 1998; Miyashita et al., 1994). Such projections could introduce motor-related modulation of sensory cortical gain, a form of dynamic normalization that adjusts responsiveness based on behavioral context (Nguyen and Kleinfeld, 2005; Seki et al., 2003). Internal, non-sensory afferents to thalamic nuclei can also depress thalamocortical synapses (CastroCAlamancos and Oldford, 2002), further shaping synchronization patterns and promoting multicluster dynamics, in which subsets of neurons synchronize without complete population entrainment (Amritkar and Rangarajan, 2009). These mechanisms could collectively explain how a heterogeneous cortical network produces the characteristically regular SWD pattern in the EEG.

A key limitation of the present study is the relatively small number of recorded neurons, reflecting the technical challenges of achieving long-term stability for juxtacellular recordings in awake animals while capturing spontaneous seizures. Nevertheless, this approach ensures that the recorded signals originate unambiguously from single neurons, which is critical when assessing neuronal heterogeneity.

Together, our findings indicate that seizures do not erase neuronal identity but modulate it in a predictable and structured manner. Neurons with distinct intrinsic and synaptic properties preserve their firing relationships across brain states, implying that the seizure state is not a chaotic departure from normal activity but a scaled reconfiguration of the pre-existing cortical network. Viewing seizures as an emergent, graded reorganization of ongoing activity rather than as a uniform cortical storm reframes our understanding of epileptic dynamics and offers new avenues for developing predictive and cell-type–specific interventions.

## Authors contributions

MLL designed research and experiments, MLL, PS, VC and NZ analyzed the data, PS and NZ conducted experiments, MLL wrote the manuscript with inputs from all authors.

## Acknowledgments

We thank Francois David and Péter Kaposvári for helpful discussions. This work was supported by the Hungarian Scientific Research Fund (Grants NN125601 and FK123831 to M.L.L.), the Hungarian Brain Research Program (grant KTIA_NAP_13-2-2014-0014), UNKP-20-5 New National Excellence Program of the Ministry for Innovation and Technology from the source of the National Research, Development and Innovation Fund to MLL , and the the Ester Floridia Neuroscience Research Foundation (grant 1502 to V.C.). MLL was a grantee of the János Bolyai Fellowship.

